# Structure-function Specialisation of the Interfascicular Matrix in the Human Achilles Tendon

**DOI:** 10.1101/2021.02.22.432199

**Authors:** Dharmesh Patel, Danae E. Zamboulis, Ewa M. Spiesz, Helen L. Birch, Peter D. Clegg, Chavaunne T. Thorpe, Hazel R.C. Screen

## Abstract

**Objective:** Tendon consists of highly aligned collagen-rich fascicles surrounded by interfascicular matrix (IFM). Some tendons act as energy stores to improve locomotion efficiency; these tendons are prone to debilitating injuries, the incidence of which increases with ageing. In equine tendons, energy storage is achieved primarily through specialisation of the IFM. However, no studies have investigated IFM structure-function specialisation in human tendons. Here, we compare the positional anterior tibialis and energy storing Achilles tendons, testing the hypothesis that the Achilles IFM has specialised composition and mechanical properties, which are lost with ageing.

**Methods:** We used a multidisciplinary combination of mechanical testing, immunolocalisation and proteomics to investigate structure-function specialisations in functionally distinct human tendons and how these are altered with ageing.

**Results:** The IFM in the energy storing Achilles tendon is more elastic and fatigue resistant than the IFM in the positional anterior tibialis tendon, with a trend towards decreased fatigue resistance with age in the Achilles IFM. With ageing, alterations occur predominantly to the proteome of the Achilles IFM.

**Conclusion:** The Achilles tendon IFM is specialised for energy storage, and changes to its proteome with ageing are likely responsible for the observed trends towards decreased fatigue resistance. Knowledge of key energy storing specialisations and their changes with ageing offers insight towards developing effective treatments for tendinopathy.

**Key messages:** *What is already known about this subject?:*

- Energy storing tendons in animals and humans are particularly prone to tendinopathy and the incidence increases with increasing age.
- Previous work in some animal models has shown that the specialisation of tendon properties for energy storage is achieved primarily through adaptation of the interfascicular matrix, with specialisation lost in ageing. However, the structural specialisations that provide the human Achilles tendon with its energy storing ability, and how these are affected by ageing, remain to be established.

*What does this study add?:*

- We demonstrate that the interfascicular matrix in the human Achilles tendon is specialised for energy storing, with increased elastic recoil and fatigue resistance, and that these specialisations are partially lost with ageing, likely due to alterations to the proteome of the interfascicular matrix.

*How might this impact on clinical practice or future developments?:*

- Short term, the specialist IFM mechanics we have demonstrated can be detected with new developments in ultrasound functional imaging, offering improved opportunities for contextual tendinopathy diagnostics. Personalised rehabilitation programmes can now be explored and designed specifically to target IFM mechanics.
- Longer term, the knowledge of key specialisations in injury prone energy storing tendons and how they are affected by ageing, offers crucial insight towards developing cell or tissue engineering treatments targeted at restoring tendon structure and function post-injury, specifically targeted at the IFM.

## INTRODUCTION

The Achilles tendon has two functions; transferring muscle-generated forces to move the skeleton, and storing energy during locomotion. Efficient energy storage requires the ability to stretch and recoil with each stride, and the mechanical properties of energy storing tendons are specialised to increase locomotory efficiency,[1,2]. The Achilles tendon is exposed to high stresses and strains during use, and has a much lower safety factor than most primarily positional tendons, such as the anterior tibialis tendon,[3,4], making it prone to age-related tendinopathy,[5-7]. While fundamental tendon structure is well defined, the structural specialisations that provide the Achilles with the mechanical properties required for energy storage remain unclear. This information is crucial to develop novel, targeted treatments that recover Achilles function post-injury.

All tendons have the same general structure, comprised of aligned, collagen-rich fascicles bound by the interfascicular matrix (IFM); a looser, less organised matrix also referred to as the endotenon,[8,9]. Our previous studies using the equine model have identified the importance of the IFM for tendon function, with structural and compositional specialisations within the IFM of the energy storing superficial digital flexor tendon (SDFT) resulting in a highly extensible and fatigue resistant material which likely translates to greater extensibility and fatigue resistance within the whole tendon,[10-14]. However, these specialisations become compromised with ageing, likely contributing to the age-related increased risk of injury to energy storing tendons in horses,[12,15,16].

While structure-function relationships in equine tendon are now well established, little work has been undertaken to elucidate structure-function relationships in human tendon or how these alter with ageing. Indeed, to the authors’ knowledge, only one study has directly compared functionally distinct human tendons, demonstrating that the Achilles has a lower elastic modulus than the anterior tibialis tendon, which is accompanied by several compositional differences,[17]. While studies have assessed the mechanical properties of the Achilles tendon *in vivo*,[1,18], or obtained healthy and/or diseased Achilles tissue via biopsy or during surgery,[19-21], the limited amount of tissue available with these approaches restricts the subsequent analysis that can be performed. Further, no studies have compared subunits of functionally distinct human tendons to establish their role in tendon structure-function relationships.

The aim of this study was therefore to compare the composition and mechanical properties of the human Achilles and anterior tibialis tendons and their subunits, and identify any age-related alterations. We hypothesise that the IFM in the energy storing Achilles tendon has specialised composition and mechanical properties, and these specialisations are lost with ageing.

## METHODS

### Patient and public involvement

Patients were not involved in design or conduct of this fundamental mechanistic study.

### Sample collection and processing

Collection, storage and use of human tendon was approved by the Human Tissue Authority (HTA; REC number: 14/NE/0154). Achilles and anterior tibialis tendons were collected, either post-mortem from the Centre for Comparative and Clinical Anatomy, University of Bristol (HTA licence: 12135), or the Newcastle Surgical Training Centre, Freeman Hospital, Newcastle upon Tyne (HTA licence: 12148), or as surgical waste from limbs amputated for tumour treatment at the Royal National Orthopaedic Hospital, Stanmore (UCL/UCLH Biobank for Studying Health and Disease; HTA license: 12055; R&D approval from UCL/UCLH/RF JRO (Ref: 11/0464)). Donors were divided into two age groups (n=9/group); middle-aged: 31–58 years, mean 47.6 years (3 female, 6 male); old aged: 72–94 years, mean 84.8 years (4 female, 5 male). Paired Achilles and anterior tibialis tendons were processed <24 hours post-mortem by quartering longitudinally, and either frozen for mechanical analysis or prepared for proteomic and histological analysis as described in supplementary methods,[12,22].

All donor tendons underwent mechanical characterisation of IFM and fascicles. Owing to the challenges associated with gripping short specimens for mechanical testing, and time required to laser-capture tissue for proteomic analysis, only 5 paired tendons from each age group underwent quarter tendon testing, histologic and proteomic analysis. See supplementary table 1 for donor and analysis details.

### Mechanical characterisation of tendon, fascicles and IFM

#### Tendon quasi-static mechanical properties

Immediately before testing, one quarter from each tendon was thawed and its cross-sectional area measured,[23]. Samples were preconditioned then pulled to failure. See supplementary methods for details of testing parameters.

#### Fascicle and IFM quasi-static mechanical properties

Samples for determination of fascicle and IFM quasi-static properties were dissected and prepared from one of the tendon quarters as previously described,[10,24]. Samples were preconditioned then pulled to failure. See supplementary methods for details of dissection and testing parameters.

#### Calculation of viscoelastic and failure properties

Force and extension data were recorded at 100Hz. Hysteresis and stress relaxation were calculated during preconditioning. For tendons and fascicles, maximum modulus, failure strain and stress were calculated during the pull-to-failure test, and for IFM samples, maximum stiffness, and force and extension at yield and failure were calculated as previously described,[11].

#### Fascicle and IFM fatigue properties

Samples for determination of fascicle and IFM fatigue properties were dissected from one of the tendon quarters and fatigue properties measured by subjecting samples to cyclic loading until failure as described previously,[13,25], with some modifications. See supplementary methods for details of testing parameters. Force and displacement data were collected at 100Hz, with cycles to failure and secondary creep rate calculated for each sample.

#### Statistical Analysis

A general linear mixed model was applied to assess differences between tendon type and with ageing, using tendon type and age as crossed factors and donor as a nested random effect factor (R, v3.6.1; www.r-project.org). Shapiro-Wilk tests indicated non-normal distribution of the data, hence data transformations using box-cox transformations were used.

### Protein Immunolocalisation

Immunohistochemistry was used to localise decorin, fibromodulin, lubricin and versican, as described previously,[22]. See Supplementary Table 2 for antibodies and blocking conditions. Elastin distribution was assessed by elastic van Gieson’s staining.

### Quantification of elastin content

The elastin content of Achilles and anterior tibialis tendons from middle-aged and old donors (n=3/group) was quantified using the FASTIN™ Elastin Assay (Biocolor, UK) as previously described,[14], and differences between tendon types assessed using paired t-tests.

### Mass Spectrometry Analyses

#### Laser-Capture Microdissection

Laser-capture microdissection of IFM and fascicles from tendon cryosections was performed as previously described,[12], with a haematoxylin staining step (1 min.) to visualise cell nuclei.

#### Protein Extraction and mass spectrometry analyses

Protein extraction and mass spectrometry analysis of laser-captured samples was performed as described previously,[12], using an Ultimate 3000 Nano system (Dionex/Thermo Fisher Scientific) coupled to a Q-Exactive Quadrupole-Orbitrap mass spectrometer (Thermo-Scientific).

#### Peptide Identification and Quantification

Fascicle and IFM proteins were identified, searching against the UniHuman reviewed database (http://www.uniprot.org/proteomes/) using parameters and filters as described previously,[26] (Peaks® 8.5 PTM software; Bioinformatics Solutions, Canada). Label-free quantification was performed separately for IFM and fascicles from each tendon and age group (Peaks® 8.5 PTM software). Protein abundances were normalised for collected laser-capture area and differentially abundant proteins between groups identified using fold change ≥2 and *p*<0.05 (PEAKS-adjusted p-values).The mass spectrometry data have been deposited to the ProteomeXchange Consortium (http://proteomecentral.proteomexchange.org) via the PRIDE partner repository,[27] with the dataset identifier PXD018212.

#### Gene Ontology and Network Analysis

The dataset of differentially abundant proteins between groups were classified for cell compartment association with Ingenuity Pathway Analysis (IPA, Qiagen) and for matrisome categories using The Matrisome Project database,[28]. Protein pathway analysis for differentially abundant proteins was performed in IPA.

## RESULTS

### Tendon, fascicle and IFM mechanical properties

There were no differences in tendon mechanical properties between age groups or tendon types, except for maximum modulus and hysteresis, which were significantly greater in the anterior tibialis tendon (Fig. 1). However, notable variability between donors was evident. At the fascicle level, ultimate tensile stress, hysteresis and stress relaxation were significantly greater in fascicles from the anterior tibialis tendon compared to those from the Achilles, however, no changes in fascicle mechanical properties were evident with ageing. Similarly, significantly greater hysteresis and force relaxation were evident in the IFM of the anterior tibialis compared to the Achilles, but no changes occurred with ageing. There were also no differences in IFM extension or force at the yield point, defined as the point at which maximum stiffness was reached and indicating the limit of elastic behaviour, between tendon types or age groups (Supplementary Fig. 1).

**Figure 1.**
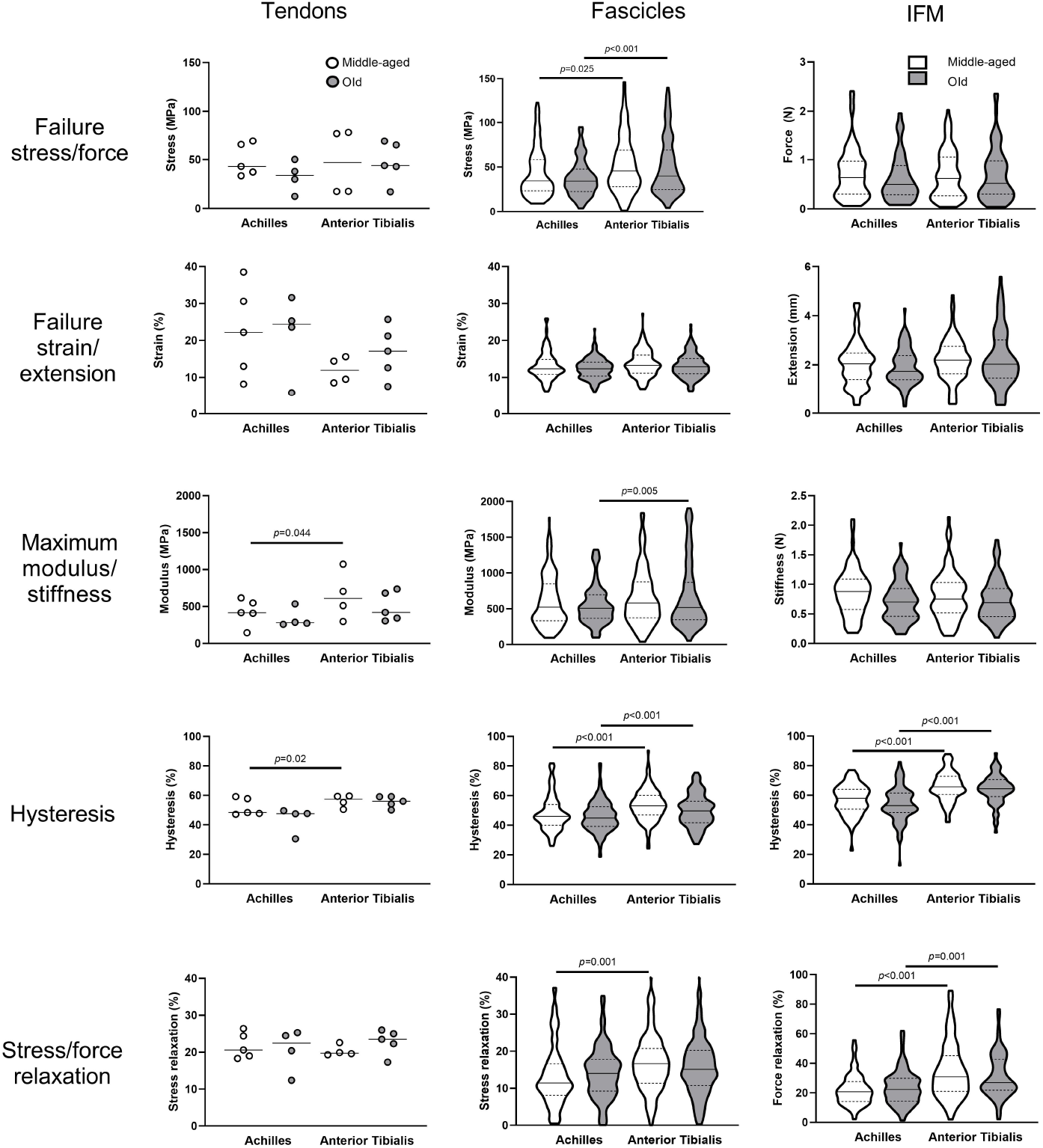
Failure and viscoelastic properties of Achilles and anterior tibialis tendons and their subunits from middle-aged and old donors. Due to limited sample numbers, individual data points are plotted for tendon tests (solid line denotes median). Distribution of fascicle and IFM data is shown by violin plots (solid line denotes median, dashed lines indicate the interquartile range and width corresponds to frequency of data points).

### Fascicle and IFM fatigue properties

Fascicle fatigue resistance did not differ between tendon types or with ageing. By contrast, the IFM in the Achilles was more fatigue resistant than that in the anterior tibialis tendon, characterised by a significantly greater number of cycles to failure (p<0.001) and lower rate of secondary creep (p<0.001). These differences were lost with ageing, due to indications of reduced fatigue resistance in the ageing Achilles IFM, seen in a trend towards a decrease in number of cycles to failure (p=0.09; Fig. 2).

**Figure 2.**
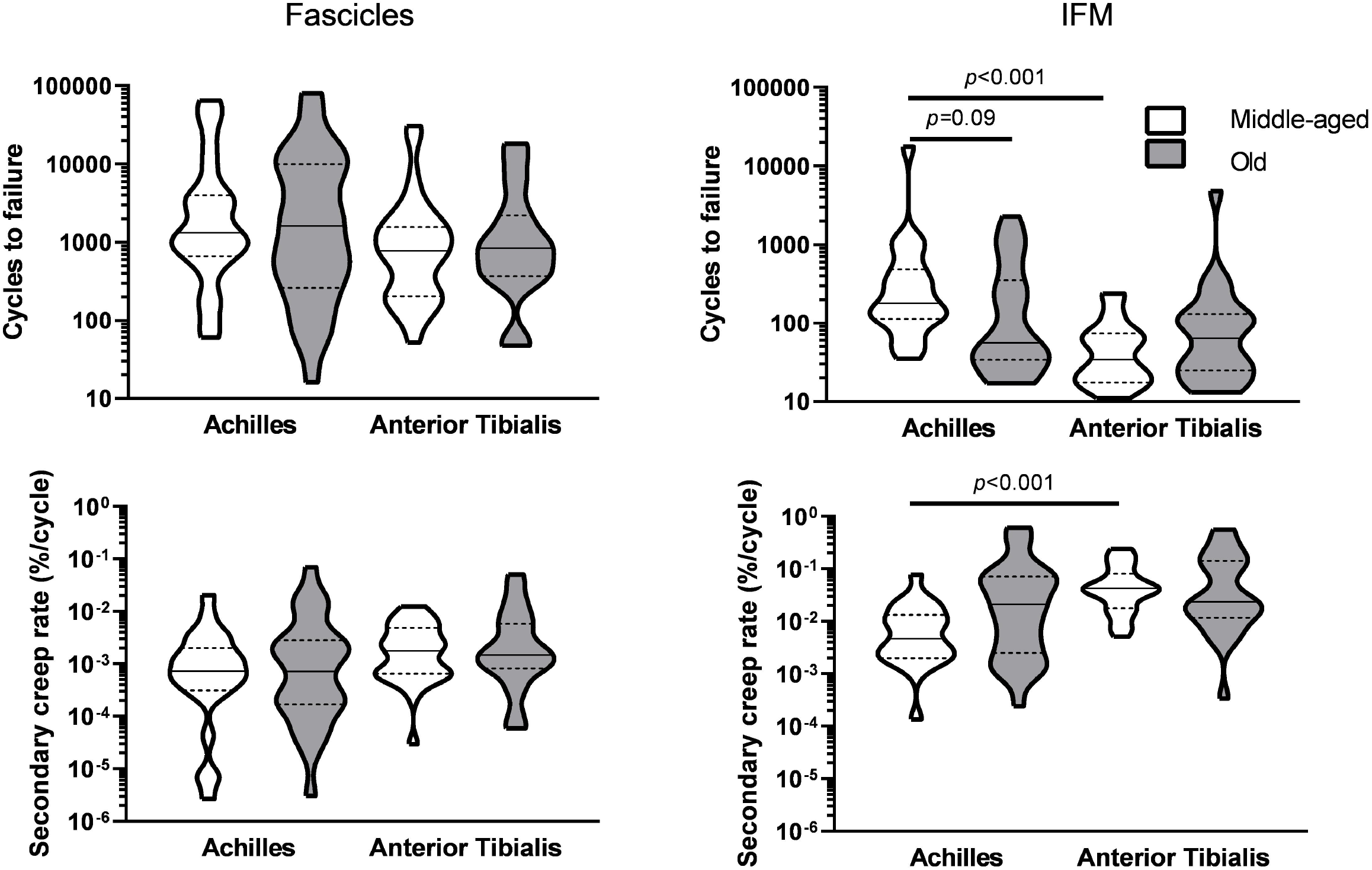
Fatigue properties of fascicles and IFM in Achilles and anterior tibialis tendons. Data are plotted on a log_10_ scale. Distribution of data is shown by violin plots (solid line denotes median, dashed lines indicate the interquartile range and width corresponds to frequency of data points).

### Protein Immunolocalisation

Lubricin, versican and elastin localised to the IFM in both tendon types. Fibromodulin staining was greater in the fascicles, with little or no staining in the IFM. Decorin was present throughout the extracellular matrix (ECM) in both tendon types (Fig. 3).

**Figure 3.**
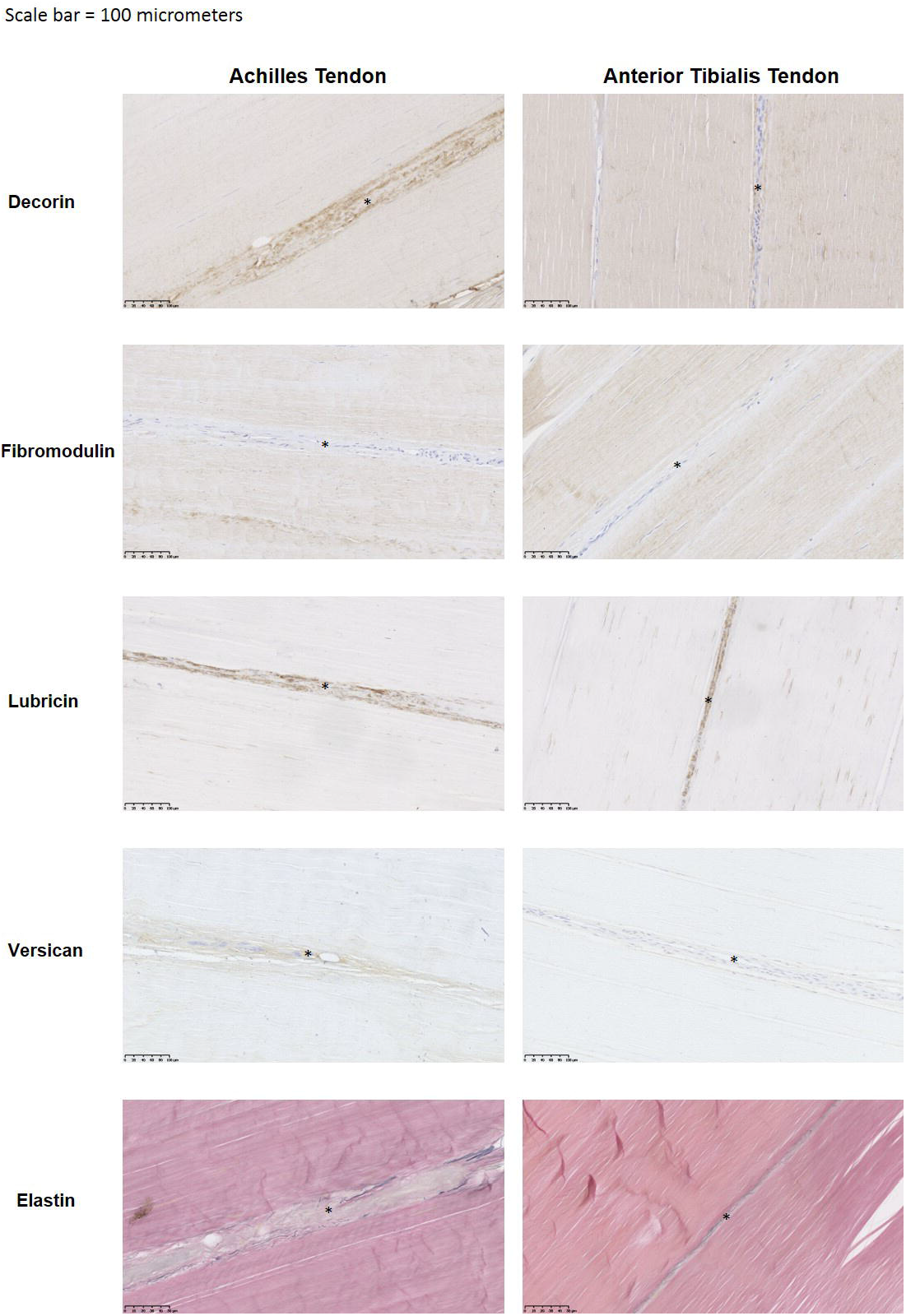
Localisation of proteins in the Achilles and anterior tibialis tendons. Representative immunohistochemical images showing distribution of tendon proteoglycans (brown) and elastin (black) in the Achilles and anterior tibialis tendons from middle-aged donors. IFM is indicated by *. Scale bar = 100 µm.

### Tendon elastin content

Elastin content in the Achilles from middle-aged and old donors was 2.4±1.6% and 1.05±0.4% respectively. Elastin content in the anterior tibialis was 1.7±0.9% in middle-aged, and 2.1±0.6% in old tendon. See supplementary figure 2 for individual data. There was a trend towards a lower elastin content in the old Achilles compared to the old anterior tibialis tendon, but this did not reach statistical significance (p=0.07).

### Protein identification and ontology

Overall, more proteins were identified in the IFM than in fascicles, and a greater proportion of those identified in fascicles were classified as ECM or ECM-related proteins (Table 1).

**Table 1.** Protein number in each of the tendon compartments as identified by LC-MS/MS and categorisation of matrisome-associated proteins. Numbers in brackets indicate percentage of total protein number.

|  | Fascicular Matrix |  |  |  | Interfascicular Matrix |  |  |  |
| --- | --- | --- | --- | --- | --- | --- | --- | --- |
|  | Achilles |  | Anterior tibialis |  | Achilles |  | Anterior tibialis |  |
|  | Middle | Old | Middle | Old | Middle | Old | Middle | Old |
| **Total Protein Number** | **152** | **152** | **148** | **153** | **265** | **259** | **243** | **211** |
| **Number of ECM Proteins** | **54**<br>**(35.5%)** | **69**<br>**(45.4%)** | **58**<br>**(39.2%)** | **53**<br>**(34.6%)** | **77**<br>**(29.1%)** | **81**<br>**(31.3%)** | **82**<br>**(33.7%)** | **74**<br>**(35.1%)** |
| ECM Glycoproteins | 16<br>(10.5%) | 22<br>(14.5%) | 17<br>(11.5%) | 16<br>(10.5%) | 28<br>(10.6%) | 31<br>(12.0%) | 31<br>(12.8%) | 29<br>(13.7%) |
| Collagens | 15<br>(9.9%) | 16<br>(10.5%) | 15<br>(10.1%) | 15<br>(9.8%) | 20<br>(7.5%) | 18<br>(6.9%) | 17<br>(7.0%) | 17<br>(8.1%) |
| Proteoglycans | 9<br>(5.9%) | 10<br>(6.6%) | 9<br>(6.1%) | 8<br>(5.2%) | 11<br>(4.2%) | 11<br>(4.2%) | 10<br>(4.1%) | 10<br>(4.7%) |
| ECM Affiliated Proteins | 4<br>(2.6%) | 6<br>(3.9%) | 6<br>(4.1%) | 5<br>(3.3%) | 9<br>(3.4%) | 8<br>(3.1%) | 9<br>(3.7%) | 7<br>(3.3%) |
| ECM Regulators | 8<br>(5.3%) | 13<br>(8.6%) | 10<br>(6.8%) | 8<br>(5.2%) | 6<br>(2.3%) | 11<br>(4.2%) | 13<br>(5.3%) | 9<br>(4.3%) |
| Secreted Factors | 2<br>(1.3%) | 2<br>(1.3%) | 1<br>(0.7%) | 1<br>(0.7%) | 3<br>(1.1%) | 2<br>(0.8%) | 2<br>(0.8%) | 2<br>(0.9%) |

#### Differences in protein abundance between tendon types and age groups

More proteins were identified as differentially abundant in the IFM than in fascicles, with most changes occurring in the old Achilles (Fig. 4&5). Many of these proteins were ECM or ECM-associated, with a predominance of proteoglycans and glycoproteins. By contrast, fewer alterations in protein abundance were observed in fascicles, and the majority of those that did change were either collagens, or were not ECM-related. Radar plots indicate that in the IFM, changes in collagen abundance were consistent for different collagen types, whereas differences in proteoglycan/glycoprotein abundance differed with protein type (Fig. 4). A similar pattern was seen for collagens in fascicles as in the IFM, with greater abundance of several collagens in the old Achilles (Fig. 5).

**Figure 4.**
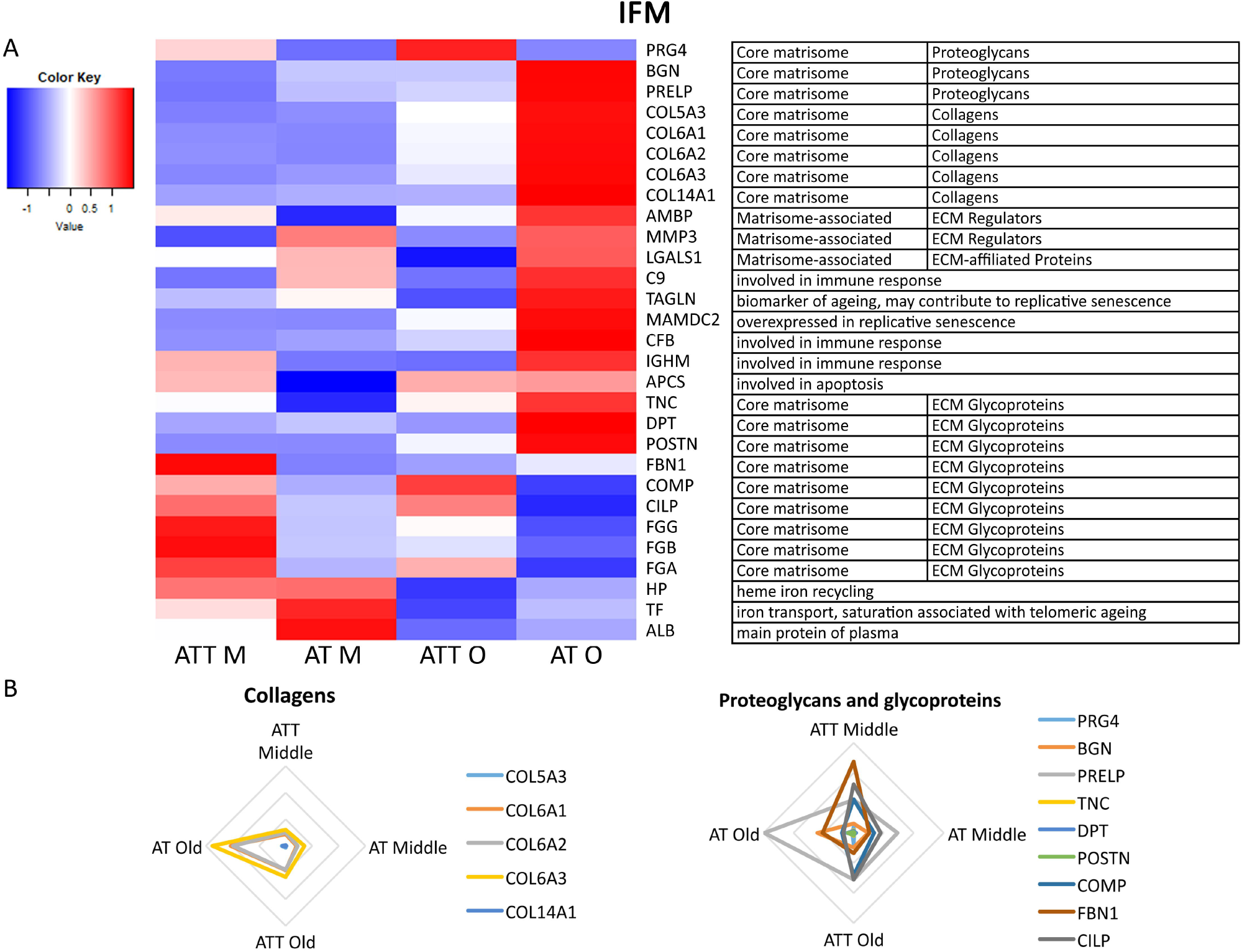
Most changes in the IFM protein abundance are observed in the old Achilles tendon. (A) Heatmap of differentially abundant proteins in the IFM middle-aged anterior tibialis (ATT M) and Achilles tendon (AT M), and old anterior tibialis tendon (ATT O) and Achilles tendon (AT O) (p<0.05, fold change ≥2). Heatmap colour scale ranges from blue to white to red with blue representing lower abundance and red higher abundance. MatrisomeDB was used to assign protein classifications. (B) Radar plots of collagens, proteoglycans and glycoproteins that showed differential abundance with age or tendon type in the IFM (p<0.05, fold change ≥2).

**Figure 5.**
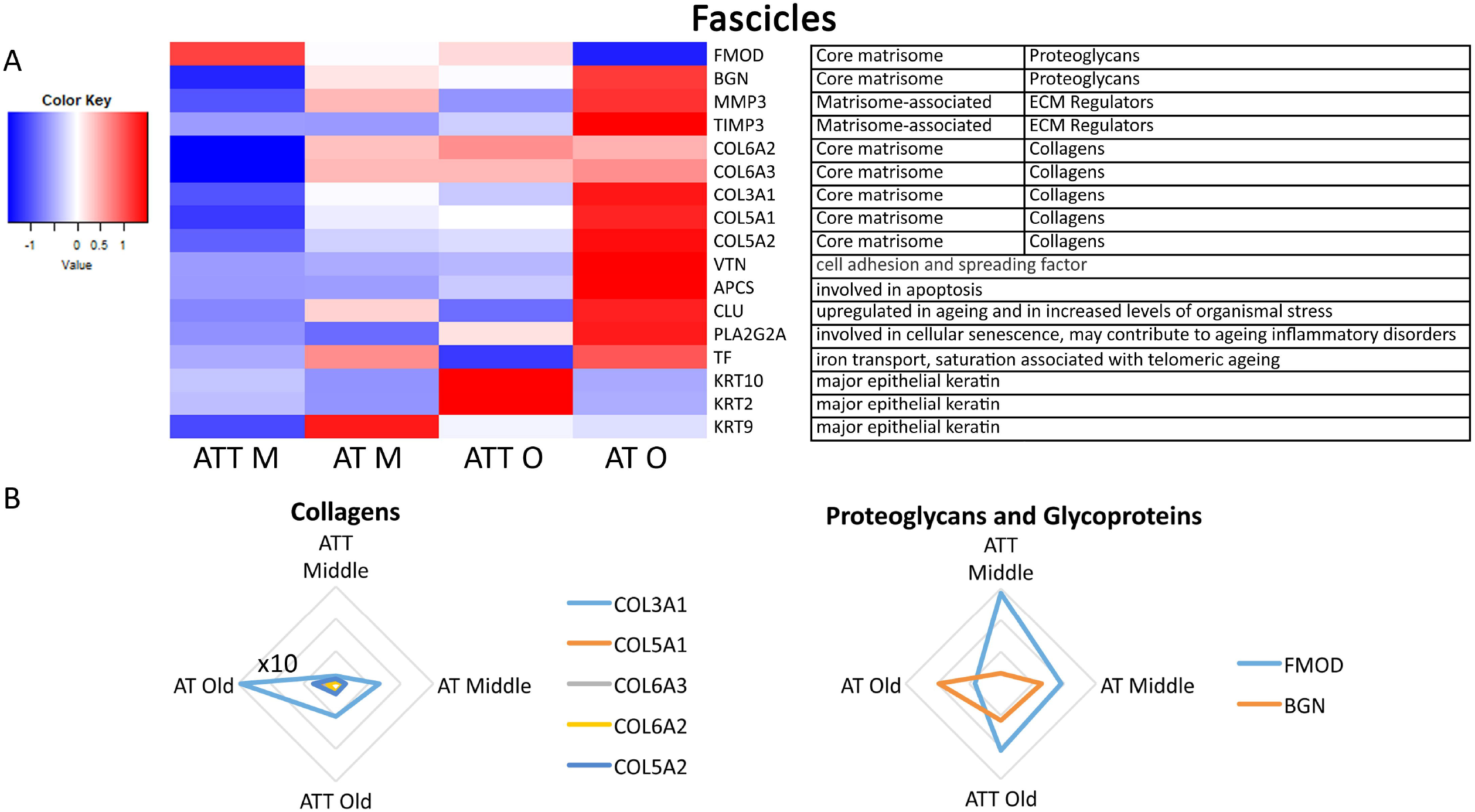
Most changes in fascicle protein abundance are observed in the old Achilles tendon. (A) Heatmap of differentially abundant proteins in the fascicles of middle-agedanterior tibialis (ATT M) and Achilles tendon (AT M), and old anterior tibialis tendon (ATT O) and Achilles tendon (AT O) (p<0.05, fold change ≥2). Heatmap colour scale ranges from blue to white to red with blue representing lower abundance and red higher abundance. MatrisomeDB was used to assign protein classifications. (B) Radar plots of collagens, proteoglycans and glycoproteins that showed differential abundance with age or tendon type in fascicles (p<0.05, fold change ≥2).

#### Pathway analysis

Potential upstream regulators were identified using IPA. TGF-β1 was predicted to be activated in the old Achilles, and inhibited in the anterior tibialis tendon, in both the IFM and fascicles (Fig. 6).

**Figure 6.**
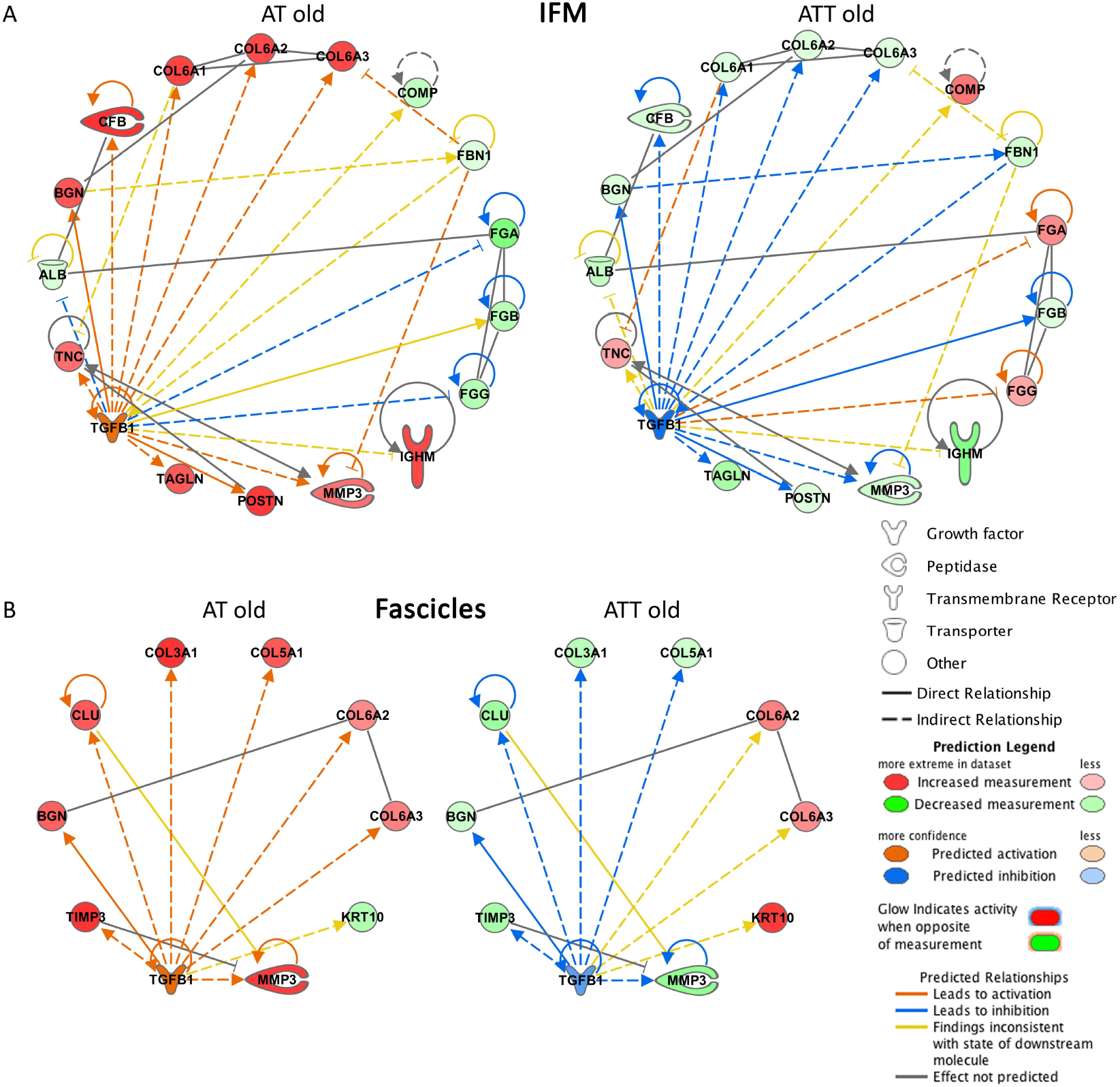
Pathway analysis identified TGF-β1 as an upstream regulator. TGF-β1 is predicted to be activated in the Achilles tendon, but inhibited in the anterior tibialis tendon from aged donors. IPA networks for TGFB1 in the IFM (A) and fascicles (B) of the Achilles tendon and anterior tibialis tendon of old donors.

## DISCUSSION

The results support our hypothesis, demonstrating specialisation of the IFM in the Achilles tendon, which is more elastic and fatigue resistant than the IFM in the anterior tibialis tendon. In contrast to our hypothesis, few changes in mechanical properties were observed with ageing, although there was a trend towards decreased fatigue resistance with ageing in the Achilles IFM only. Proteomic analysis revealed a more complex proteome in the IFM, with age-related alterations in protein abundance predominantly occurring in the Achilles IFM.

Overall, the results we present here are similar to those reported previously in functionally distinct equine tendons, with greater elasticity in energy storing compared to positional tendons and their subunits,[10,11]. However, we observed fewer differences between human tendons than seen previously in equine tendon, in which whole tendons and fascicles also show several differences in mechanical properties between tendon types,[10,13]. This may simply arise from the lower numbers of available human samples and high sample variability, but may also be associated with the unmatched age ranges of human and equine samples, or with differences in energy storing function, as the Achilles is a less extreme energy store than the highly specialised equine SDFT,[29,30].

We were not able to obtain tendons from donors under 30 years old, so we are comparing middle-aged and old, rather than young and old, providing less age contrast than in our previous studies of equine tendon. It is possible that specialisation of the Achilles was already diminishing in the middle-aged group, as we observed previously in equine tendon,[15], so fewer age-related alterations in mechanical properties were evident. Indeed, it is well established that Achilles tendinopathy is most prevalent in the fourth decade of life,[7,31], which may well result from this diminishing specialisation coinciding with continual or increasing usage as individuals take up new sporting activities.

However, it is also apparent that there are large variations in tendon mechanical properties between individuals, which may mask differences between tendon types or with ageing. The source of this variation is uncertain, but it should be noted that we had a mixed-sex population, and minimal information about the health, exercise or injury status of donors. Tendons with any macroscopic signs of degeneration were excluded from all analyses. However, whilst some donors did have documented diabetes, in others information of any systemic diseases were lacking. While the sex of each donor was known, limited sample numbers means it was not possible to establish if any variability arose from sex-related differences at baseline or with ageing, as previously reported,[32]. Despite these limitations, it is notable that we still identified significant compositional and mechanical differences between energy storing Achilles and positional anterior tibialis tendons.

Few studies have investigated the mechanical or structural properties of functionally distinct human tendons and their subunits. Those that have, typically analyse a single tendon type to explore limited mechanical or compositional aspects. The IFM of the human Achilles tendon has received little attention previously, with data supporting the results we present here, demonstrating localisation of lubricin to the IFM,[33], and identifying capacity for interfascicular sliding,[34]. Indeed, a recent modelling study indicated that sliding of tendon subunits, enabled by IFM, is necessary to accurately predict tendon viscoelasticity and failure,[35]. However, in contrast to our previous findings in functionally distinct equine tendons,[10,11,15], we did not identify any differences in interfascicular sliding between tendon types or age groups, as measured by IFM extension at the point of maximum stiffness. While the capacity for interfascicular sliding does not appear to differ between tendon types, our results demonstrate enhanced elasticity and fatigue resistance in the Achilles IFM, specialisations that likely contribute to efficient energy storage in this tendon.

Histological analysis confirmed that the IFM is rich in proteoglycans, particularly lubricin and decorin, and also elastin, as seen previously in tendons from other species,[22,36,37]. Mass spectrometry allowed a comprehensive characterisation of the IFM and fascicular proteomes and comparison between tendon types and age groups, revealing a greater complexity in the IFM proteome in both tendon types, with almost double the number of glycoproteins identified in this region compared to the FM, supporting previous findings in equine tendon,[12]. The protein profile identified is also similar to that detected previously in the whole Achilles tendon, with many collagens and proteoglycans present,[38]. Elastin detection by mass spectrometry requires the inclusion of an elastase digestion step,[38] which was not possible with our samples due to the limited volumes collected by laser capture. However, by combining quantitative assays to determine whole tendon elastin content, and immunolocalisation to identify its spatial arrangement, we were able to establish that elastin was localised to the IFM, with a trend towards lower elastin content in the old Achilles compared to the old anterior tibialis tendon. Previous research demonstrates a decline in elastin content in the energy storing equine SDFT with ageing,[14]. In our samples, a particularly large individual variation in elastin content in the middle-aged Achilles tendon was evident (supplementary figure 1), which may mask a decline in elastin content in this tendon with ageing. Previous studies have demonstrated highly variable energy storage capacity between the Achilles tendons of middle-aged individuals,[39], which also indicates significant individual variability in human tendons with ageing.

While no significant changes in tendon or subunit quasi-static mechanics were identified with age, the superior fatigue resistance of the Achilles IFM was lost with ageing, suggesting some effect of ageing on the Achilles IFM specifically. Indeed, proteomic analysis revealed that the majority of changes in protein abundance were measured in the IFM from old Achilles tendons relative to the other groups. Many of these changes were ECM-related, suggesting dysregulation of homeostasis which may be responsible for the loss of superior fatigue resistance observed. This is in contrast to previous transcriptomic analysis of the ageing Achilles, which showed little changes in ECM proteins at the gene level,[40]. It may be that the changes in ECM protein abundance we observed are post-transcriptionally regulated, or that separate analysis of IFM and fascicles allows detection of differences that are not apparent when the tendon is analysed as a whole. Indeed, very few proteins changed in abundance in both fascicles and IFM with ageing, instead ageing changes occurred predominantly in the IFM, suggesting differential age-related regulation of protein homeostasis across fascicles and IFM. Of interest, we measured increased abundance of collagen VI in the Achilles IFM with ageing. Collagen VI is overexpressed in several fibrotic diseases, and collagen VI-null mice exhibit improved cardiac structure and function post-myocardial infarction,[41,42].

TGF-β signalling was predicted to be activated in the Achilles tendon from old donors. TGF-β signalling is essential for tendon development, and is expressed predominantly within the IFM of developing and adult tendons,[43-45]. Indeed, our recent work demonstrates upregulation of TGF-β in the IFM of the energy storing equine SDFT upon commencement of loading during development,[26]. However, TGF-β signalling has also been shown to be dysregulated in several age-associated diseases, including atherosclerosis, neurodegenerative diseases and arthritis, and is upregulated in tendon injury,[46,47]. In cartilage, TGF-β switches from a protective to a detrimental role with ageing, which is associated with osteoarthritis development,[48]. It is possible that dysregulation of TGF-β signalling in the old Achilles drives the changes in the IFM proteome reported here.

## Conclusions

In this study, we demonstrate specialisation of the IFM in the energy storing Achilles tendon, with greater elasticity and fatigue resistance than in the positional anterior tibialis tendon. Further, we identify age-related alterations in the IFM proteome of the Achilles tendon which is likely related to the loss of fatigue resistance observed. These changes may contribute to the increased risk of Achilles tendinopathy with ageing, and provide information crucial for developing improved tendinopathy diagnostics and IFM-targeted therapeutics.

## Author Contributions

DP: Investigation, formal analysis, writing – review & editing; DEZ: Investigation, formal analysis, visualisation, writing – review & editing; EMS: Investigation, writing – review & editing; HLB: Conceptualisation, funding acquisition, writing – review & editing; PDC: Conceptualisation, funding acquisition, supervision, writing – review & editing; CTT: Conceptualisation, funding acquisition, writing – original draft; HRCS: Conceptualisation, supervision, funding acquisition, writing – review & editing.

## Acknowledgements

The authors would like to thank Dr Deborah Simpson for her assistance with mass spectrometry experiments and data analysis

