## Supplementary Information for "Structure-function Specialisation of the Interfascicular Matrix in the Human Achilles Tendon"

| **Age** | **Male or Female** | **Use** | | | | |
| --- | --- | --- | --- | --- | --- | --- |
|  |  | **Histology** | **Proteomics** | **FM/IFM mechanics** | **FM/IFM fatigue** | **1/4 tendon mechanics** |
| 31 | F | x | x | x | x | x |
| 37 | F |  |  | x |  | x |
| 43 | F |  |  | x |  | x |
| 46 | M | x | x | x | x | x |
| 49 | M | x | x | x | x | x |
| 51 | M |  |  | x |  | x |
| 56 | M | x | x | x | x | x |
| 57 | M | x | x | x | x | x |
| 58 | M |  |  | x |  | x |
| 72 | M |  |  | x |  | x |
| 79 | M |  |  | x |  | x |
| 80 | M |  |  | x |  | x |
| 83 | F |  |  | x |  | x |
| 87 | F | x | x | x | x | x |
| 87 | F | x | x | x | x | x |
| 93 | F | x | x | x | x | x |
| 93 | M | x | x | x | x | x |
| 94 | M | x | x | x | x | x |

**Supplementary Table 1. Details of donors and tests performed on samples from each donor.**

| **Primary Antibody** | **Supplier** | **Dilution** | **Secondary Antibody** | **Supplier** | **Dilution** | **Blocking conditions** |
| --- | --- | --- | --- | --- | --- | --- |
| Decorin | Atlas Antibodies; HPA003315; rabbit IgG | 1:50 | Peroxidase-goat anti rabbit IgG | Sigma; A6154 | 1:50 | 20% goat serum in TBS-Tween |
| Fibromodulin | Provided by Prof. Roughley, McGill University; rabbit IgG | 1:50 | Peroxidase-goat anti rabbit IgG | Sigma;  A6154 | 1:50 | 20% goat serum in TBS-Tween |
| Versican | Abcam; ab19345; rabbit IgG | 1:50 | Peroxidase-goat anti rabbit IgG | Sigma; A6154 | 1:50 | 20% goat serum in TBS-Tween |
| Lubricin | MDB biosciences; 1042015; mouse IgG | 1:50 | Peroxidase-goat anti mouse IgG | Sigma; A4416 | 1:50 | 20% goat serum in TBS-Tween |

**Supplementary Table 2. Details of primary and secondary antibodies used for immunohistochemistry of matrix proteins.** Negative controls were included where primary antibodies were omitted; no staining was visible in any of these samples.

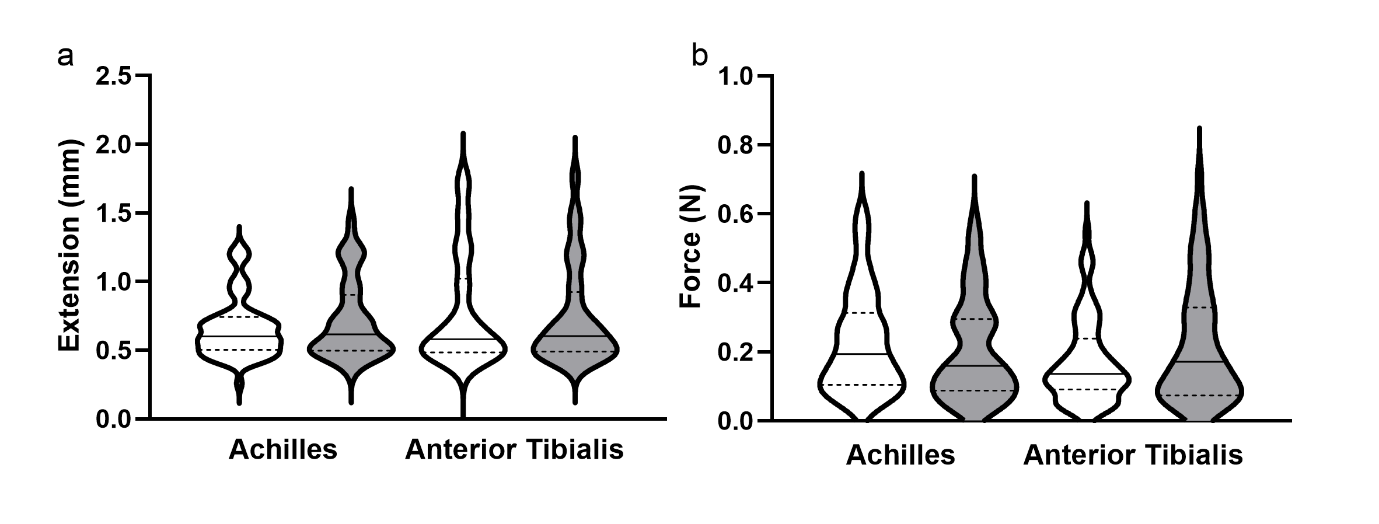

**Supplementary Figure 1. Extension and force at the yield point in IFM samples.** Distribution of data is shown by violin plots (solid line denotes median, dashed lines indicate the interquartile range and width corresponds to frequency of data points).

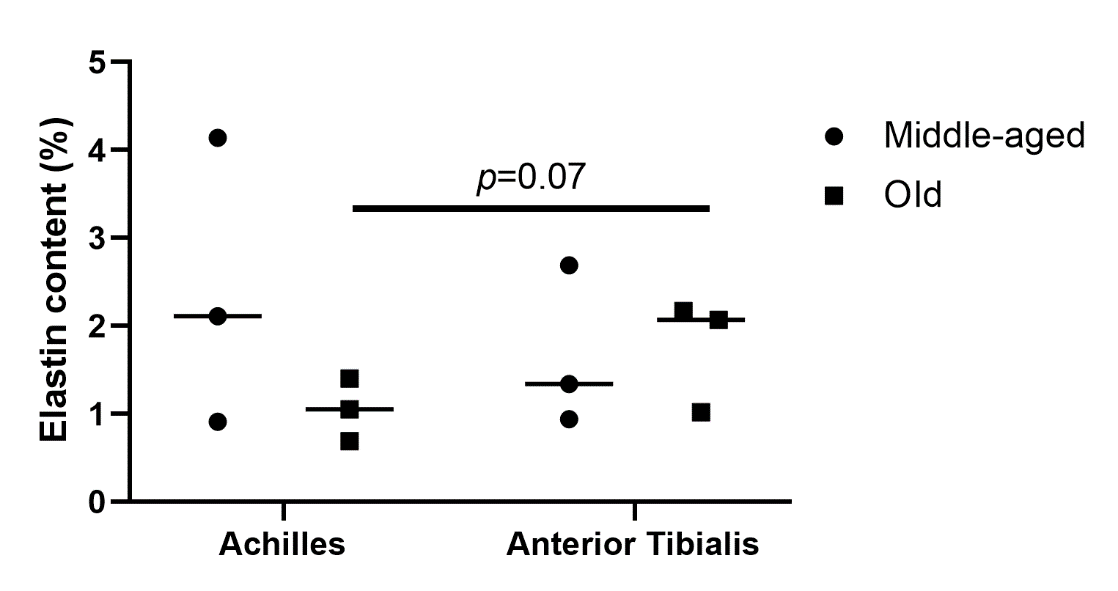

**Supplementary Figure 2. Elastin content in Achilles and anterior tibialis tendons from middle-aged and old donors.** Lines indicate median values.
