## Supplementary Methods for "Structure-function Specialisation of the Interfascicular Matrix in the Human Achilles Tendon"

*Sample processing* - Three of the quarters from each tendon were wrapped in tissue paper dampened with phosphate buffered saline and stored at -20°C to enable future mechanical analysis. Two small segments (approx. 5 x 5 x 5 mm^3^) were taken from the remaining quarter for proteomic and histological analysis as previously described,[1, 2]. For proteomics, segments were embedded in OCT, snap frozen in hexane cooled on dry ice and stored at -80°C, and for histology, segments were fixed in 4% paraformaldehyde at 4°C for 24 hours and then wax embedded. Visual inspection of tendons during dissection did not identify any abnormalities.

*Tendon* *quasi-static mechanical properties* – Samples were clamped in a materials testing machine (Instron 5967, 30kN load cell) using custom-made cryogrips cooled with liquid CO_2_, with an average grip to grip distance of 50 mm. A preload of 20 N or 10 N was applied for Achilles tendon and anterior tibialis tendons respectively (equivalent to 2% of the predicted failure force) and the gauge length was measured as the distance between the freeze lines at either end of the tendon. Samples were preconditioned for 10 loading cycles from 0 to 2.5% strain at 1 Hz, followed by a pull to failure at 5% strain/second.

*Fascicle and IFM quasi-static mechanical properties* - fascicles, approximately 40 mm in length were dissected by cutting longitudinally through the tendon and fascicle diameter measured using a laser micrometer ,[3]. IFM samples were prepared from 2 adjoining fascicles by cutting through the opposing ends of the fascicle pairs such that 10 mm of IFM was left intact between the two fascicles, enabling the IFM to be tested in shear by pulling the opposing fascicle ends in a uniaxial manner. Specimens were secured in a materials testing device (Electropuls E1000, 250N load cell, Instron) using pneumatic grips with a grip to grip distance of 20 mm, and a preload of 0.02 N applied. Specimens were preconditioned to 2.5% strain (fascicles) or 0.25 mm (IFM) (approximately 25% of failure strain/extension) for 10 cycles at 1 Hz and then pulled to failure at 5% strain/second ,[4].

*Fascicle and IFM fatigue properties* - Specimens were secured in custom-made testing chambers with a grip to grip distance of 10 mm and hydration maintained by filling the chamber with DMEM ,[5]. Chambers were gripped in a materials test machine (Electroforce 5500, 225 N load cell, TA Instruments) and a preload of 0.02 N applied to the specimens. Having previously demonstrated the failure strains are more consistent than the failure stresses between samples, a single loading cycle was used to identify the appropriate peak load for cyclic creep tests. For fascicle fatigue loading, one loading cycle to 60% of the predicted fascicle failure strain was applied, and the maximum load recorded was then applied to fascicles in a cyclic manner at 1 Hz frequency until sample failure. A similar procedure was used for IFM fatigue tests, except that a loading cycle to 50% predicted failure extension was used to determine the load to apply during the subsequent fatigue test.
